# Localizing the Sources of Diffusion Mediating Structure-Function Mapping Using Graph Diffusion Wavelets

**DOI:** 10.1101/2024.09.07.611772

**Authors:** Chirag Jain, Sravanthi Upadrasta Naga Sita, Avinash Sharma, Bapi Raju Surampudi

## Abstract

The intricate link between brain functional connectivity (FC) and structural connectivity (SC) is explored through models performing diffusion on SC to derive FC, using varied methodologies from single to multiple graph diffusion kernels. However, existing studies have not correlated diffusion scales with specific brain regions of interest (RoIs), limiting the applicability of graph diffusion. We propose a novel approach using graph heat diffusion wavelets to learn the appropriate diffusion scale for each RoI to accurately estimate the SC-FC mapping. Using the open HCP dataset, we achieve an average Pearson’s correlation value of 0.833, surpassing the state-of-the-art methods for prediction of FC. It is important to note that the proposed architecture is entirely linear, computationally efficient, and notably demonstrates the power-law distribution of diffusion scales. Our results show that the bilateral frontal pole, by virtue of it having large diffusion scale, forms a large community structure. The finding is in line with the current literature on the role of the frontal pole in resting-state networks. Overall, the results underscore the potential of graph diffusion wavelet framework for understanding how the brain structure leads to functional connectivity.

**AUTHOR SUMMARY:** In the network diffusion paradigm for brain structure-to-function mapping, we noticed limitations such as manually decided diffusion scales and the absence of RoI-level analysis. We addressed this problem by independently developing the graph diffusion wavelets having multiscale and multiresolution property. Each brain region is associated with a diffusion scale that defines the extent of spatial communication. Using graph diffusion wavelets, we are able to predict the functional connectome with state-of-the-art (SoTA) results. We observe that the diffusion scales follow a power-law degree distribution, which is indicative of a scale-free process in the brain. The frontal pole is a dominant member of the various resting-state networks, and our model is able to associate higher diffusion scales to this region. The graph diffusion wavelet model is a novel method which not only excels in downstream task but also provides insights into the structure-function relation.

## INTRODUCTION

The physical properties of biological entities are a key to understanding and predicting various functional capabilities. The principle of “structure determines function” has been extremely influential in molecular biology, biochemistry and physical sciences (Alberts et al., 2002; Michael, 2021). Neuroscientists have tested this dogma on the brain and have observed that brain structural connectivity largely determines its functional connectivity (Damoiseaux & Greicius, 2009; Sporns, 2013). The structural connectivity (SC) is derived from diffusion tensor imaging (dMRI) which captures the white matter tracts (myelinated axons) internally connecting the gray matter of the cortex. These tracts are obtained by a tractography algorithm and then with the help of an atlas that divides the cortex in multiple regions, we can determine the strength of structural connectivity between cortical regions. The static functional connectivity (FC) is derived from functional MRI (fMRI) that is based on the blood oxygen level dependent activity (BOLD signal) sampled across the duration of resting state epoch. The voxel-wise BOLD signal, again with the help of the same atlas, can be converted to region-of-interest (RoI)-specific signal which represents local aggregate of oxygen concentration at different time points. Further, we can calculate the Pearson’s correlation between the BOLD signal of each of the RoIs to get connectivity strength based on correlation in their activityZ.-Q. Liu, Betzel, and Misic (2022). Various fMRI studies have shown that brain RoIs interact to form functional networks when involved in visual (Lowe, Dzemidzic, Lurito, Mathews, & Phillips, 2000), language, (Hampson, Peterson, Skudlarski, Gatenby, & Gore, 2002) or working memory (Hampson, Driesen, Skudlarski, Gore, & Constable, 2006) tasks (Skudlarski et al., 2008) and also in resting-state (Biswal, Zerrin Yetkin, Haughton, & Hyde, 1995).

Mapping structural to functional connectivity is essential to understand the intricate network dynamics of the brain and its implications for various neurological disorders. This mapping allows researchers to discover biomarkers associated with diseases such as Autism Spectrum Disorder (Chen et al., 2021), Alzheimer’s (Patow et al., 2023), etc., by analyzing deviations between predicted FC derived from SC and empirical FC data. Such studies facilitate the creation of reliable computational models of brain networks that can be tested against specific perturbations associated with a variety of diseases, ultimately improving our understanding of the neural circuitry responsible for behavioral dysfunctions. By bridging the gap between structural and functional perspectives, researchers can better elucidate the mechanisms underlying cognitive processes and pathologies, paving the way for targeted therapeutic strategies and interventions in clinical settings.

Modeling the brain as a graph through network neuroscience allows for a comprehensive analysis of its structure and function by representing brain regions as nodes and anatomical connections as edges (Bassett & Sporns, 2017; Bassett, Zurn, & Gold, 2018). This approach, applicable from micro-to macro-scales, facilitates the creation of connectomes that link structural and functional domains, revealing meaningful properties of brain networks (Papo & Buldú, 2024). Using graph theory techniques, we can gain insight into the complexities of brain connectivity and dynamics, enhancing our understanding of both healthy and pathological states.

One of the main goals of connectomics is to understand how and to what extent brain structure influences its function. For over a decade, various studies relying on network organization have tried to establish a relationship between structure and function (Bullmore & Sporns, 2009; Honey et al., 2009; Honey, Thivierge, & Sporns, 2010). This network-based approach enables the use of graph theory methods such as random walk (Rosenthal et al., 2018) and spectral graph theory methods like graph Fourier transform (Griffa, Amico, Liégeois, Van De Ville, & Preti, 2022), which are based on the principle of graph heat diffusion. Abdelnour et al. (Abdelnour, Voss, & Raj, 2014) proposed that brain functional states (FC) can be predicted using a single heat diffusion kernel, a method that proved very successful and prompted further studies to rely on variants of such network diffusion models. Later, Surampudi et al. (Surampudi et al., 2019, 2018) demonstrated that FC can be better explained using a linear combination of heat diffusion kernels. With the advent of geometric deep learning (GDL), methods like graph convolutional neural network (GCN) (Kipf & Welling, 2017) and graph transformer network (GTN) (Yun, Jeong, Kim, Kang, & Kim, 2019) based encoder-decoder models were used for finding better mappings (Ji, Deslauriers-Gauthier, & Deriche, 2021).

However, GCN-based learning methods are susceptible to the problem of over-smoothing where all the node features converge to the same representation in the latent space, more so in a fully connected brain graph. Further, since GDL methods lacked explainability, they were eventually integrated with graph diffusion methods (Oota, Yadav, Dash, Bapi, & Sharma, 2022, 2024) that rely on attention mechanism for combining the heat kernels. Although incorporating attention leads to somewhat superior results compared to previous methods, they suffer from high computational complexity.

In this paper, we address the above issues by using the graph diffusion wavelets which localize the diffusion kernel, enabling each brain region to have a unique diffusion scale. Implicit multi-resolution nature of wavelets allow us to have RoI-level information of interaction in the brain. The proposed method is entirely linear, has lower space-time complexity, and at the same time performs better than the previous state-of-the-art (SoTA) methods.

## MATERIALS AND METHODS

### Theory

The graph wavelets, with a rich history rooted in physics and mathematics, are explored in this section, where we explain graph diffusion, its relation to computational neuroscience, and introduce the wavelet concept.

#### Graph Diffusion

The heat diffusion equation is a partial differential equation which models the process of heat flow in space. For example, consider a metal rod, one end of which is heated. The heat will flow to another, relatively cooler end, and this process can be modelled using the diffusion equation (1).

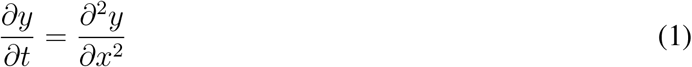

However, the graph is a discrete mathematical entity and therefore the equation is discretized, leading to the formulation of a Laplacian operator *L*. The Laplacian operator encodes spatial information of the graph, and the eigenfunctions of graph Laplacian form the basis set for graph Fourier transform. The solution **y(t) = h**_**t**_**(y) = e**^**−sL**^**y(0)** is a heat diffusion kernel which operates on **y** to determine the extent of diffusion after some time *t* parameterized by diffusion scale *s*. The matrix exponential can be computed as follows:

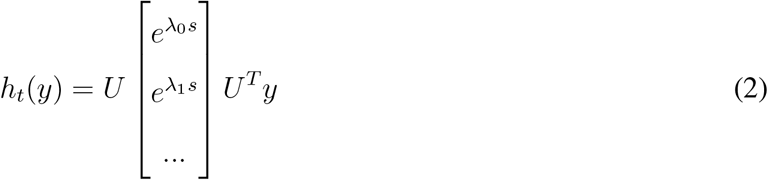

where *U* is the eigenvector matrix and (*λ*_0_ … *λ*_*N*−1_) are the eigenvalues of Laplacian L. The diffusion equation is prevalent in many branches of science (Stewart, 1999) and technology (Sharma, Horaud, Cech, & Boyer, 2011; Sharp, Attaiki, Crane, & Ovsjanikov, 2022).

#### Neural basis of heat diffusion

The Wilson-Cowan equation (Wilson & Cowan, 1972) is one of the most successful models of interaction between populations of excitatory and inhibitory neurons. It is an instance of Reaction-Diffusion (RD) equations wherein the activity diffuses over space, and the diffusion is guided by the connection strength between elements of that space. *Turing mechanism* for RD systems states that the spatial modes of the Laplacian of a system are instrumental in understanding the perturbation of that system (Kondo & Miura, 2010; Van Gorder, 2021). Atasoy *et al*. (Atasoy, Donnelly, & Pearson, 2016) emphasizes the use of Laplacian operator across all physical systems and specifically its utility in modelling the RD systems to eventually characterize the “connectome harmonics”. These connectome harmonics are the stationary waves associated with the graph, which can resolve the graph activity based on its spatial frequencies. The state of such system at any time is given by the heat kernel given in equation 2.

#### Graph Wavelets

In signal processing, wavelets are used to localize the signal in the time domain. This allows exploration of properties of the signal for specified time period instead of the entire time series data. Time-frequency plots are one such example which is extensively used in EEG signal processing Cohen (2014). Similar to time domain, we can have wavelets in the spatial domain which can localize the spatial data (like a graph). Wavelets can be applied to graphs to capture the local neighborhood of specific nodes (Coifman & Maggioni, 2006; Gavish, Nadler, & Coifman, 2010; Hammond, Vandergheynst, & Gribonval, 2011). Wavelets at different locations and spatial scales are formed by translating and scaling the mother wavelet (Hammond et al., 2011).

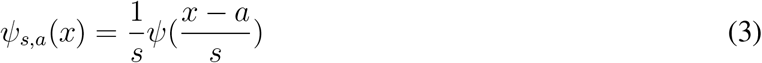

where *a* is the *a*^th^ node and *s* is the scaling factor. A graph wavelet captures the local neighborhood of a node when a unit energy is propagated from the node to the graph (Donnat, Zitnik, Hallac, & Leskovec, 2018). Moreover, nodes can operate at different diffusion scales, helping us understand the role of each node within the graph. A significant limitation of the heat kernel is its lack of spatial localization, which greatly reduces its explainability, and this can be fixed by wavelets.

In this paper, we propose the use of graph wavelets to localize heat diffusion, providing a higher resolution view of the RD process. By associating different diffusion scales with each node, we can uncover the properties of various brain Regions of Interest (RoIs) with greater precision and clarity. This approach enhances our ability to understand the intricate dynamics within the graph, like the existence of scale-free networks, offering a significant improvement over previous methods.

### Graph Wavelet Diffusion

For a brain region *i*, we write the rate equations by considering all the connected regions. The number of firing neurons in region *j* is given by *Ŝ*_*i,j*_ *x*_*j*_(*t*) where *Ŝ*_*i,j*_ is the (*i, j*)^th^ entry of unnormalized SC, representing the number of tracts between *i*^th^ and *j*^th^ regions. The net change in number of firing neurons in region *i* is considered a linear function of the number of active neurons from region *j*, which according to (Abdelnour et al., 2014) gives:

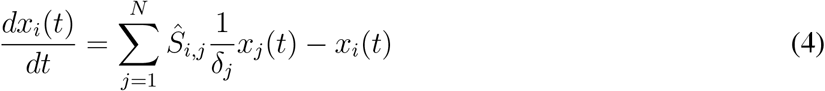

*δ*_*j*_ being the degree of node *j*.

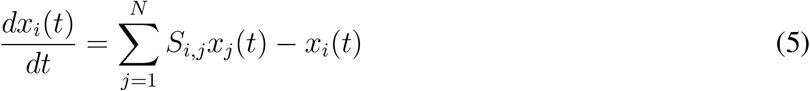

where 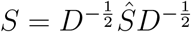 is the row normalized structural connectivity matrix. Concatenating over *i* results in:

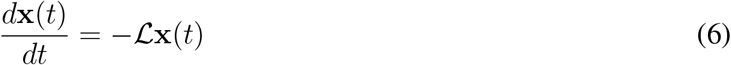

where ℒ = **I** − *S*. The solution for above equation is:

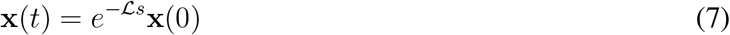

For multi-resolution and multiscale analysis, we perform dot product with an impulse function and have a unique diffusion kernel for each node.

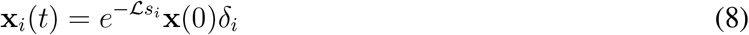

Without loss of generality, assume that **x**(0) is a unit vector and the matrix exponential can be computed through Laplacian eigen-decomposition along with a diagonal heat kernel matrix (*G*).

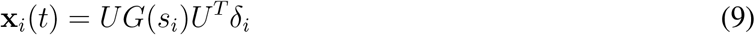

where 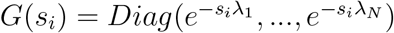. We can construct an asymmetric kernel matrix (**K**) by concatenating all the wavelets together.

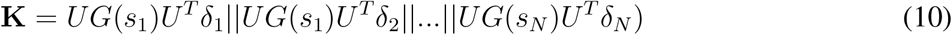

At time *t*, the configuration of any given region *i*, based on an initial configuration, represents the functional connectivity of region *i* with all other regions. To obtain FC, we concatenate the asymmetric kernel and SC and pass it through a per-vertex linear layer (see Figure 3), which enables us to learn the optimal diffusion scales. In other words, a linear layer is utilized for learning the mapping between each row of the concatenated matrix (**K**||**S**) to each row of FC. The diffusion process utilizes the neighborhood structure, while the linear layer performs mapping for individual nodes. This linear layer can be replaced by MLP by addition of an activation function, but note that our primary model is entirely linear.

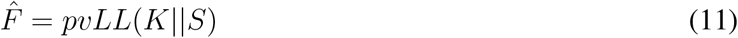

Here, *pvLL*() refers to the process whereby a linear layer (*LL*) is applied on each vertex (*pv*) of the concatenated matrix to eventually yield a predicted FC matrix 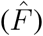. The non-linear MLP does not capture the spatial information but helps in encoding the high-frequency features (Sharp et al., 2022).

**Figure 1.**
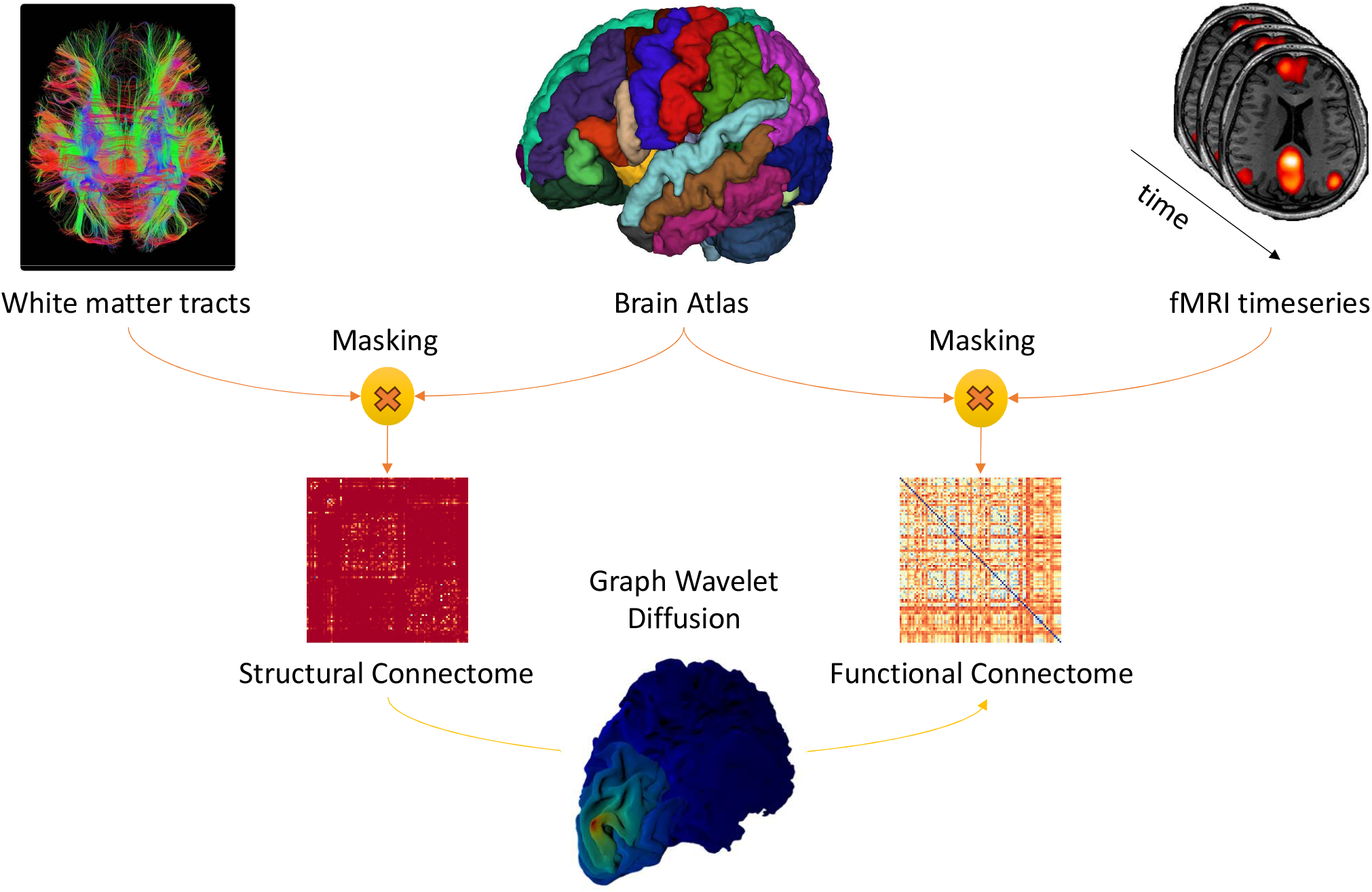
The white matter tracts are extracted from diffusion MRI and the activity is estimated using functional MRI. Both are masked using a brain atlas to obtain the structural connectivity matrix and region-wise time series data of brain activation, which is used for computing the functional connectivity matrix.

**Figure 2.**
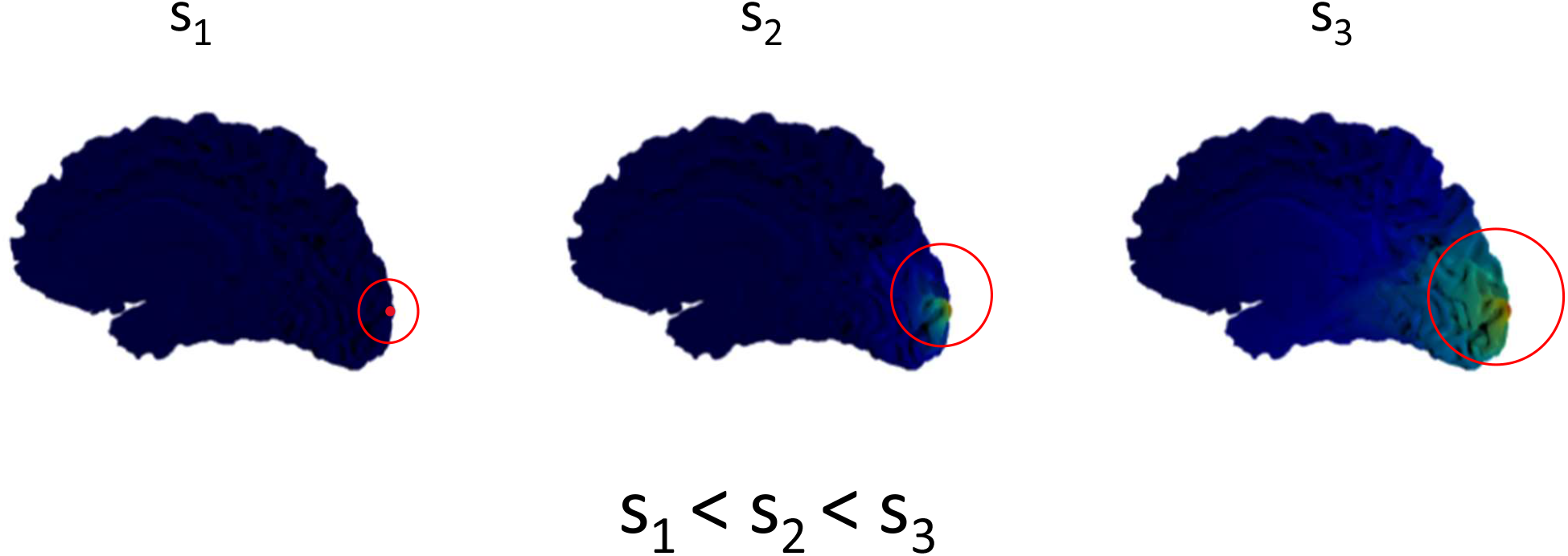
Visualization of diffusion on the brain surface mesh. We show the wavelet diffusion for 3 different scales in increasing order. Note that higher diffusion scales corresponds to bigger neighborhood.

**Figure 3.**
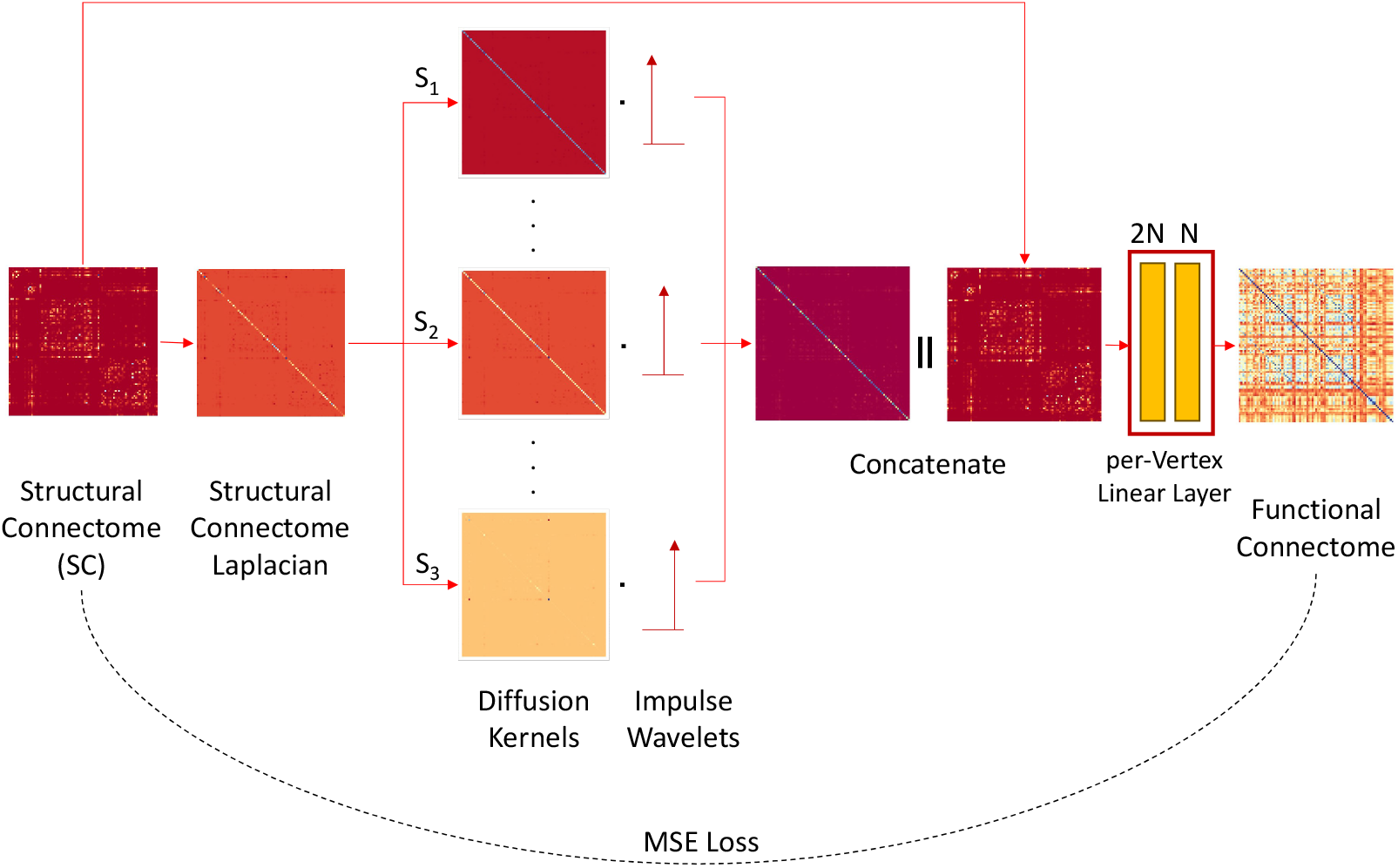
Flow of the Graph Wavelet method. 1. Calculate normalized graph Laplacian of operator (*L* = *D* − *A*), where *L, D*, and *A* are graph Laplacian, degree, and adjacency matrices, respectively. 2. Calculate heat kernel for each node as follow: 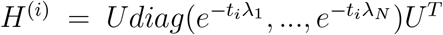 where *H*^(*i*)^ is the heat kernel for *i*^*th*^ node and *U* is the eigenvector matrix of Laplacian. 3. Extract brain region from corresponding heat kernel by localization with its dot product with impulse function. This step gives an asymmetric heat kernel which can be concatenated with SC, thus combining neighborhood information. 5. Lastly, flatten each node and map it to the ground truth FC using linear layer for each node.

### Training

In this section, we define the update rule for the diffusion scales and other aspects of training. We use the Adam optimizer (Kingma & Ba, 2015) along with a scheduler and the Frobenius norm of difference between predicted and groundtruth FC. Here, SC and FC are represented by *X* and *Y*, respectively.

Forward Pass:

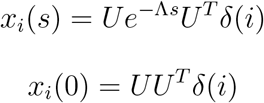

Since *x*_*i*_(0) is constant, gradients corresponding to it will be zero.

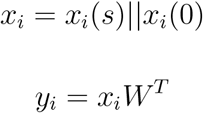

where *y*_*i*_ is the *i*^*th*^ row of FC matrix. We concatenate all the *y*_*i*_ and then calculate the loss. The loss function is defined as the Frobenius norm of difference of predicted FC [*Ŷ*] and groundtruth FC [*Y* ] (referred to as MSE loss in Figure 3). When projecting a matrix onto the space of positive semidefinite matrices (like FC), the nearest positive semidefinite matrix can be computed using the Frobenius norm (Higham, 1988).

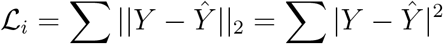

Backward Pass:

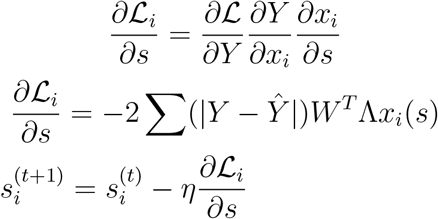

As can be seen from the update rule for the diffusion scales, the scales of RoIs are dependent on the overall functional connectivity, the linear mapping *W*, and the derivative of the wavelet at scale *s*.

### Scale Selection

The question of which scales to use always persisted in the literature, right from (Abdelnour et al., 2014) and MKL (Surampudi et al., 2018) to A-GHN (Oota et al., 2024). We solve this problem by learning the scales by relying on backpropagation learning algorithm with the MSE loss function as shown in Figure 3.

### Data

The dMRI and fMRI datasets are from the Human Connectome Project (HCP1200) dataset. We assessed the 1058 subjects provided in the HCP1200 data release (Essen et al., 2013). The data is processed by (Zhang et al., 2018) and made available publicly at the GitHub repository. They have used the Desikan et al. (2006) atlas, consisting of 68 cortical (34 from each hemisphere) and 19 subcortical brain regions. As mentioned in the repository, the 19 subcortical regions are not consistently arranged across SC and FC. We resolved this issue by reordering the FC matrix so that the correspondence between RoIs of SC and FC is maintained. This data processing step is crucial for our work, since graph wavelets are first localized and then mapped to specific nodes.

## RESULTS

We conduct rigorous experiments on our model, comparing it quantitatively with previous SoTA models, and also explain the patterns observed in learned diffusion scales. The plan experiments is as follows. We undertake comparison of performance of the proposed model with respect to SoTA models and cross validation experiments on the proposed model itself. Various ablation experiments are then conducted to investigation the impact of changing the number of eigen basis functions used for estimation of predicted FC; effect of using an MLP layer in place of a linear output layer; and finally investigate various features of the learned diffusion scales.

### 5 random runs

As the proposed model utilizes multi-resolution, multi-scale graph diffusion wavelets, we perform comparative study on a graph diffusion-based approach that does not use deep networks such as multiple kernel learning (MKL) model (Surampudi et al., 2018). Further, we chose models that do not utilize diffusion but implement geometric deep learning methods such as GCN Encoder-Decoder model (Li, Shafipour, Mateos, & Zhang, 2019) and another model that combines Graph Convolutional Network (GCN) and Graph Transformer Network (GTN) (Ji et al., 2021). Finally, we considered a model that combines graph diffusion, heat kernels and deep neural networks with attentional mechanism, namely, attention graph heat network (AGHN) model (Oota et al., 2024) for comparative analysis.

All the models underwent training and testing for a total of five times on the open HCP dataset that has 1058 subjects. For each run, the dataset was partitioned into three distinct parts: 50% was allocated for training, comprising 529 samples; 5% was designated for validation, consisting of 53 samples; and the remaining 45% was set aside for testing, encompassing 476 samples. Across these five iterations, the model’s performance was evaluated using the Pearson’s correlation coefficient (PCC) metric. The results of all the previous models were taken as reported from (Oota et al., 2024).

As can be seen from Table 1, on average, the Pearson’s correlation coefficient for the proposed (linear) model between predicted FC and empirical FC was 0.8326 *±* 0.0017, indicating a strong positive relationship between the predicted and actual values. For visual confirmation, refer to figure 4 which shows the groundtruth and predicted functional connectivity matrices for a randomly selected subject.

**Table 1:**
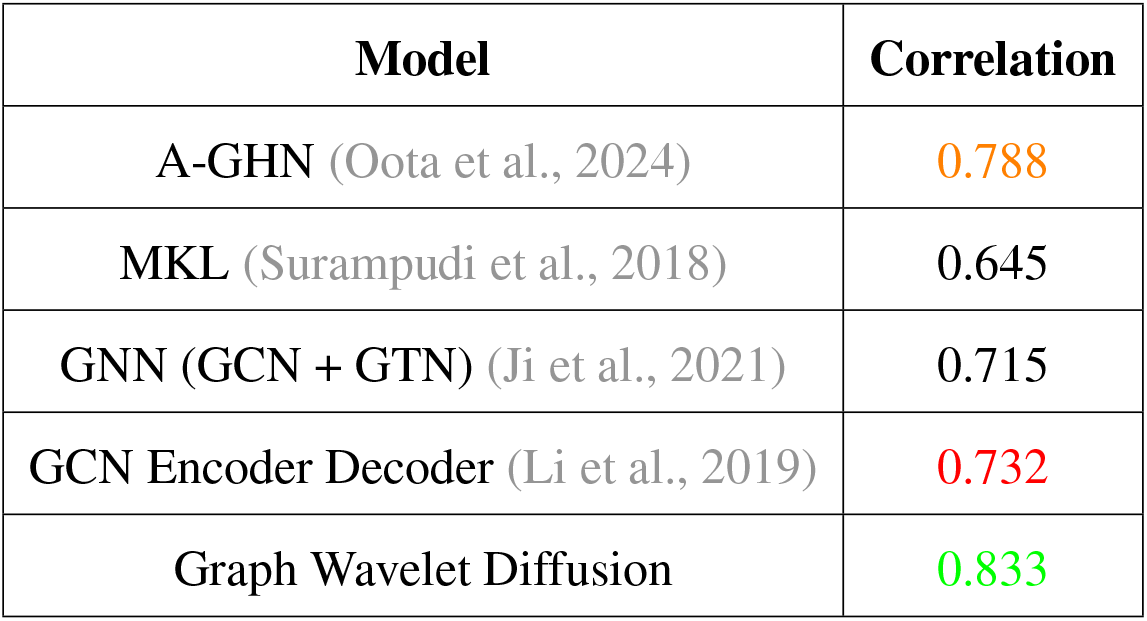
Performance comparison of graph diffusion wavelet method with previous methods.

| Model | Correlation |
| --- | --- |
| A-GHN (Oota et al., 2024) | 0.788 |
| MKL (Surampudi et al., 2018) | 0.645 |
| GNN (GCN + GTN) (Ji et al., 2021) | 0.715 |
| GCN Encoder Decoder (Li et al., 2019) | 0.732 |
| Graph Wavelet Diffusion | 0.833 |

**Figure 4.**
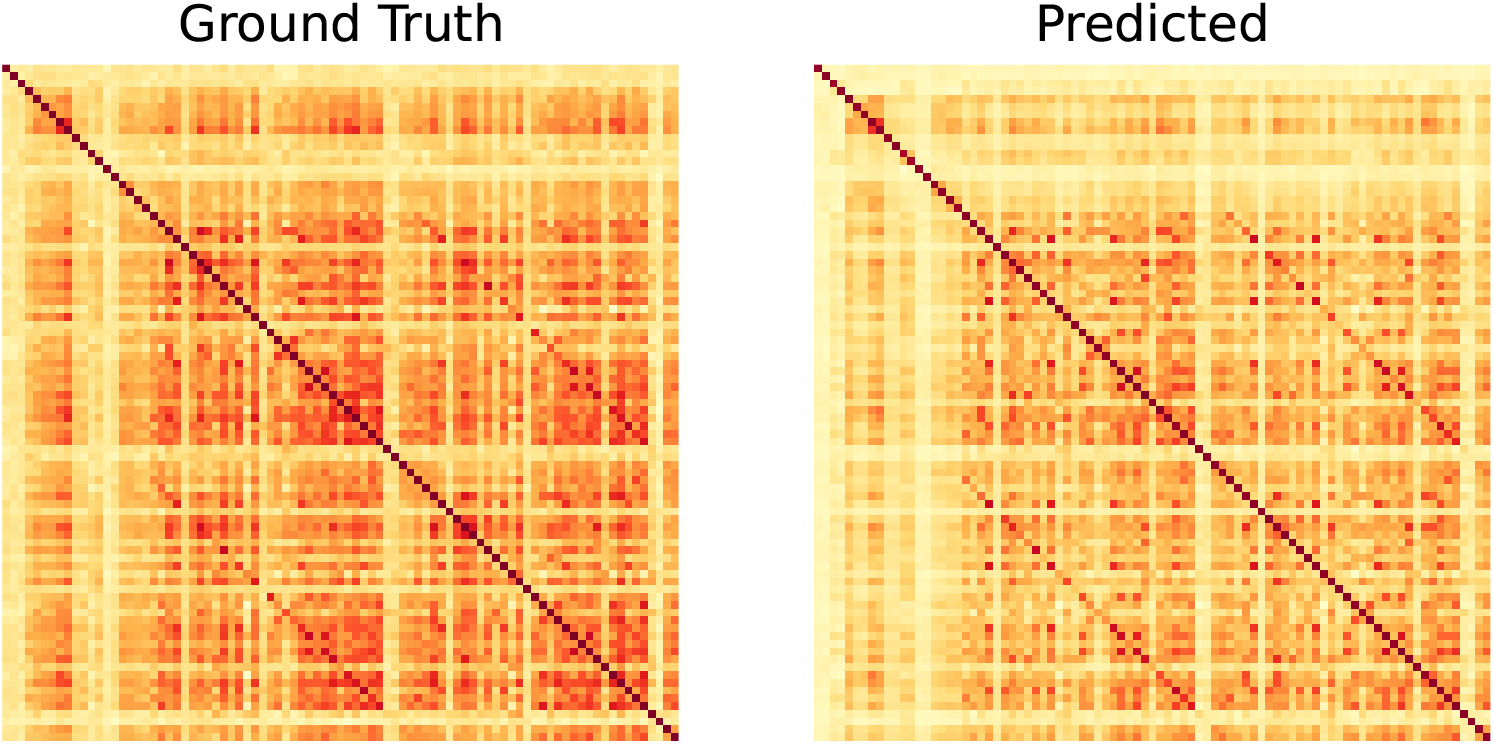
Groundtruth and predicted FC matrix presented for qualitative evaluation for a randomly chosen subject.

Overall, we achieve significantly better performance as compared to A-GHN (previous SoTA) while being around 4 times smaller in size (number of parameters) and double in training speed. These details will be discussed further in the following sections.

### 5-fold Cross Validation Experiments

In order to further investigate the impact of varying the train-test splits we conducted 5-fold cross validation (CV) experiments. The dataset was divided into 5 folds, with each fold containing around 211 subjects. For each run, the model was trained on 4 of these folds, leaving the remaining fold for testing. This cross-validation process was repeated until each fold had been used as the test set once. The performance results on cross-validation experiments are illustrated in Figure 5. Notably, the Pearson’s correlation coefficient is close to 85%, indicating a strong positive relationship between the predicted and empirical FC in those cases irrespective of the fold.

**Figure 5.**
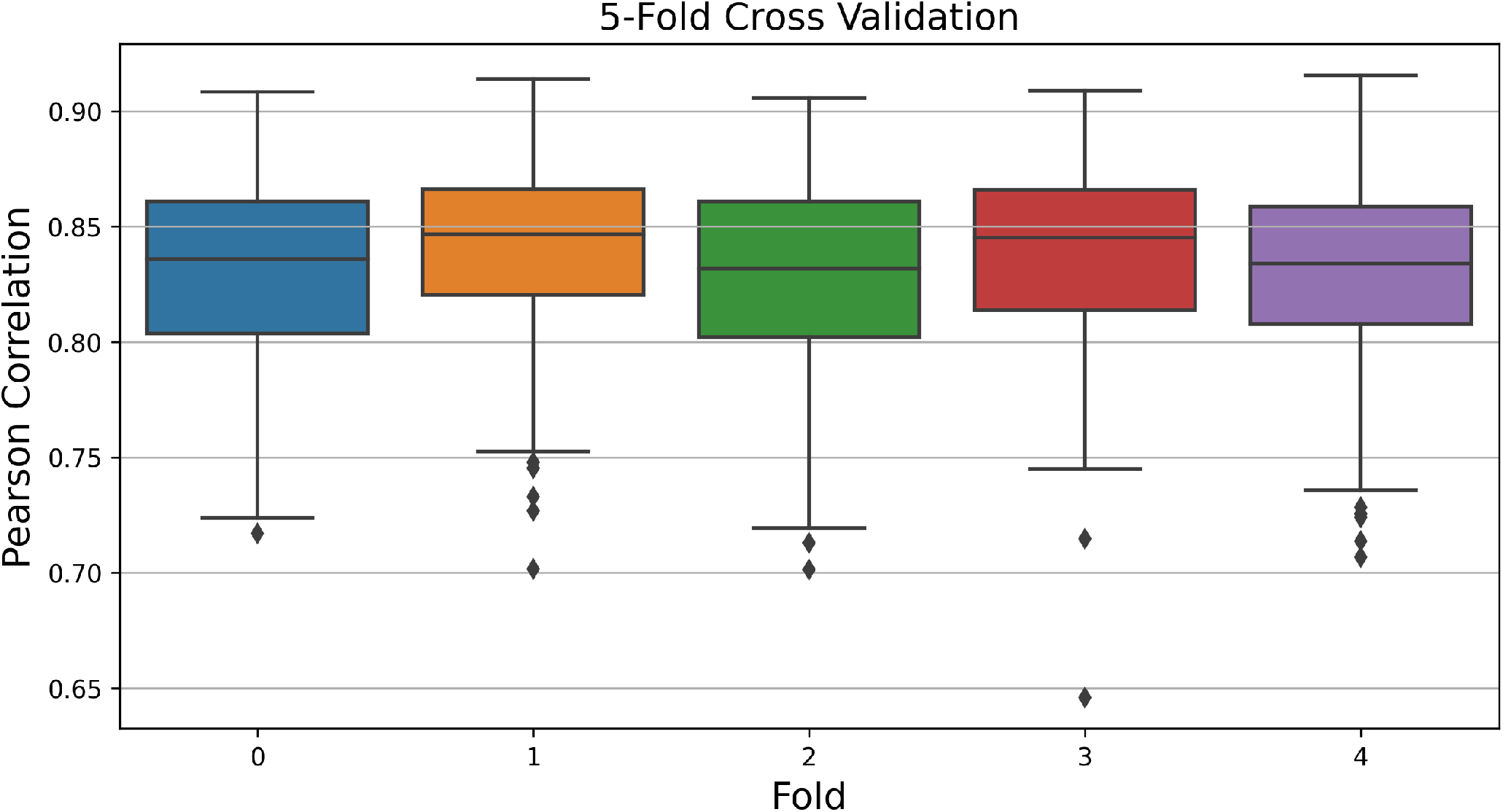
5-fold cross validation results. Each box-plot depicts results on the 210 subjects in the testing fold.

### Impact of Number of Basis Functions and MLP Layers

The eigen functions that form the basis set for the Laplacian represent spatial harmonics that can be utilized for filtering purposes. It is possible to discard the basis functions corresponding to higher spatial frequencies to evaluate their significance for FC reconstruction. Consequently, when constructing heat kernels, one can exclude the basis functions associated with high spatial frequencies rather than using the complete basis set (*U* ∈ ℝ^*N ×k*^). In Table 2, results are reported when 87, 75, and 64 low-frequency basis sets are used for computing the heat kernel. The findings indicate that higher spatial frequencies play a crucial role in the reconstruction of FC, as their exclusion leads to a noticeable decline in performance in both the PCC and MSE metrics.

**Table 2:** Performance of model for varying number of basis sets and size of the MLP hidden layer.

| k (No. Eigenvectors) | MLP | Pearson's Correlation |  |  | MSE |  |  |
| --- | --- | --- | --- | --- | --- | --- | --- |
|  |  | Train Set | Validation Set | Test Set | Train Set | Validation Set | Test Set |
| 87 | Linear | $0.8332 \pm 0.0014$ | $0.8318 \pm 0.0059$ | $0.8326 \pm 0.0017$ | $0.0220 \pm 0.0004$ | $0.0228 \pm 0.0017$ | $0.0223 \pm 0.0004$ |
| | 64 | $0.8314 \pm 0.0008$ | $0.8269 \pm 0.0016$ | $0.8283 \pm 0.0004$ | $0.0223 \pm 0.0003$ | $0.0236 \pm 0.0006$ | $0.0225 \pm 0.0005$ |
| 75 | Linear | $0.8292 \pm 0.0010$ | $0.8224 \pm 0.0079$ | $0.8267 \pm 0.0013$ | $0.0224 \pm 0.0004$ | $0.0241 \pm 0.0016$ | $0.0226 \pm 0.0004$ |
| | 64 | $0.8277 \pm 0.0010$ | $0.8237 \pm 0.0039$ | $0.8254 \pm 0.0006$ | $0.0224 \pm 0.0003$ | $0.0231 \pm 0.0020$ | $0.0228 \pm 0.0003$ |
| 64 | Linear | $0.8200 \pm 0.0006$ | $0.8133 \pm 0.0048$ | $0.8171 \pm 0.0003$ | $0.0230 \pm 0.0006$ | $0.0241 \pm 0.0017$ | $0.0233 \pm 0.0005$ |
| | 64 | $0.8237 \pm 0.0019$ | $0.8180 \pm 0.0036$ | $0.8206 \pm 0.0012$ | $0.0226 \pm 0.0007$ | $0.0237 \pm 0.0016$ | $0.0231 \pm 0.0008$ |

### Learned Diffusion Scales

We extracted the learned diffusion scale values from 10 random runs (the train-validation-test split remains as for the 5-random run experiments reported earlier). We chose a larger number of runs to get a smoother estimate of the diffusion scale value. The PCC values for the predicted FCs over the 10-run experiments remain similar to those reported earlier for the 5-run experiments. In this section, we plot the average values of diffusion scales learned over the training epochs as shown in Figure 6. Most of the nodes have low diffusion rate, while few nodes display a larger scale for optimal FC prediction. Larger scale correspond to larger neighborhood while smaller scales corresponds to small neighborhood. The Frontal Pole exhibits largest diffusion scale. Note that all the diffusion scales were initialized at 0 and the increase in scale value for certain regions is based on the learning process that happens in the downstream task of mapping SC to FC.

**Figure 6.**
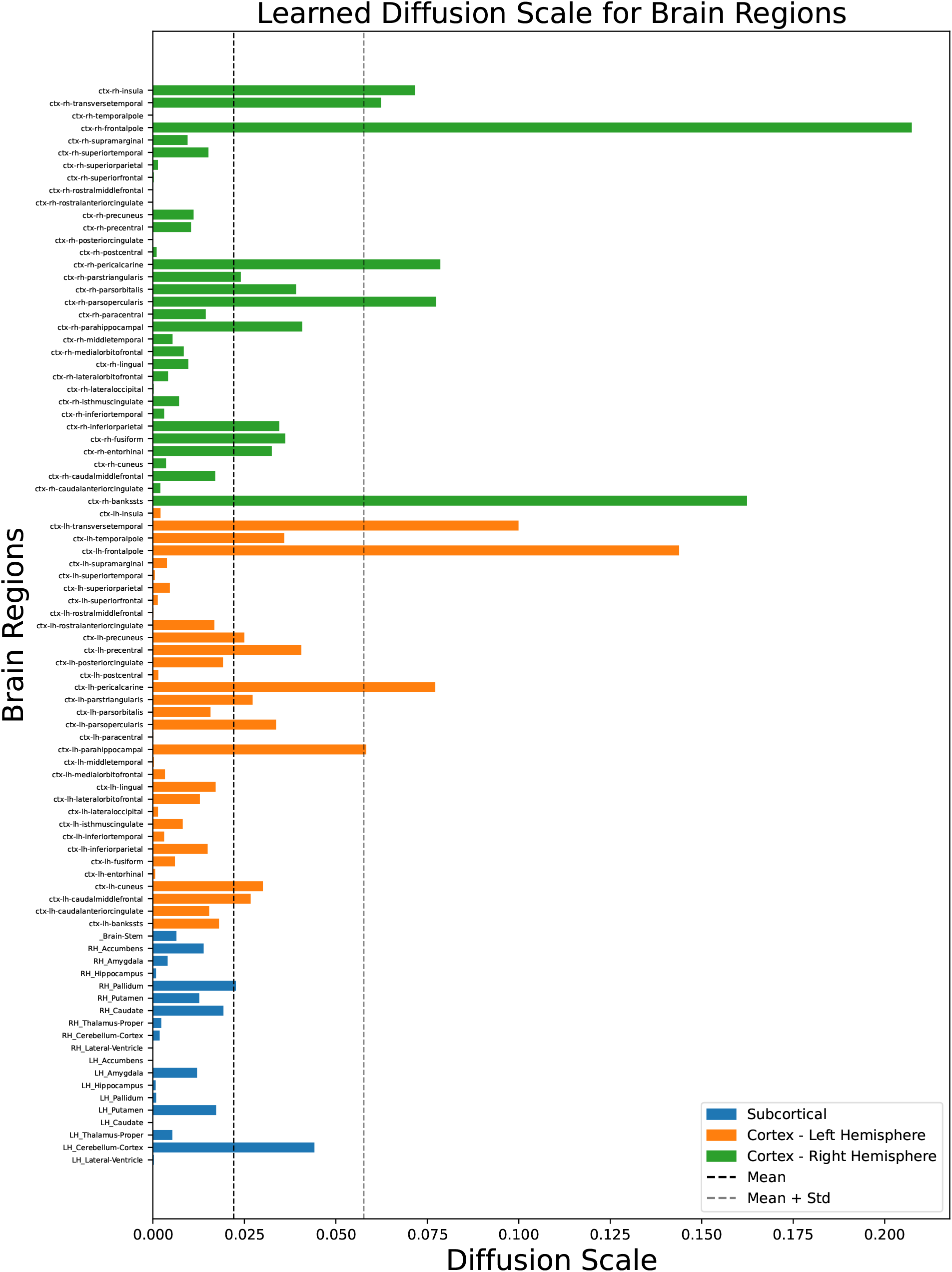
Average value of the diffusion scale (averaged over 10 runs) learned for each brain region. The labels follow the order in the SC/FC, starting from bottom to top.

### Scale-free Property

Diffusion scales correspond to the size of the neighbourhood of a node in the graph. In this sense they appear to capture similar information as node-degree in a graph. It is well-known that node-degree distribution follows power-law in scale-free networks such as brain graphs (Bassett & Bullmore, 2006). It would be interesting to see if learned diffusion scale values also conform to this property. In order to test this possibility, we plot the histogram of diffusion scale Figure 7a and try to fit the power-law and exponential curves to analyze the pattern in the diffusion scales. The diffusion scale describes the spatial extent to which a node interacts with its graph, effectively controlling the neighborhood of that node. Degree distribution in a scale-free network follows a power-law, wherein few nodes have large degree and most nodes have small degree (Barabási & Albert, 1999; Ciuciu, Varoquaux, Abry, Sadaghiani, & Kleinschmidt, 2012; He, 2011; He, Zempel, Snyder, & Raichle, 2010; Wang & Chen, 2003). The existence of scale-free property in the connectomes has been found previously, and our study corroborates such findings (Achard, Salvador, Whitcher, Suckling, & Bullmore, 2006; Bassett & Bullmore, 2006; Ciuciu et al., 2012; He, 2011; He et al., 2010; Ponce-Alvarez, Kringelbach, & Deco, 2023). We fit the data to the power-law and exponential curves and find that the sum of squared estimates of error (SSE) for power law is around half as compared to that of the exponential distribution as shown in Figure 7a. Based on Clauset, Shalizi, and Newman (2009), we estimate the power-law exponent as 2.04 and the corresponding log-log plot can be seen in Figure 7b.

**Figure 7.**
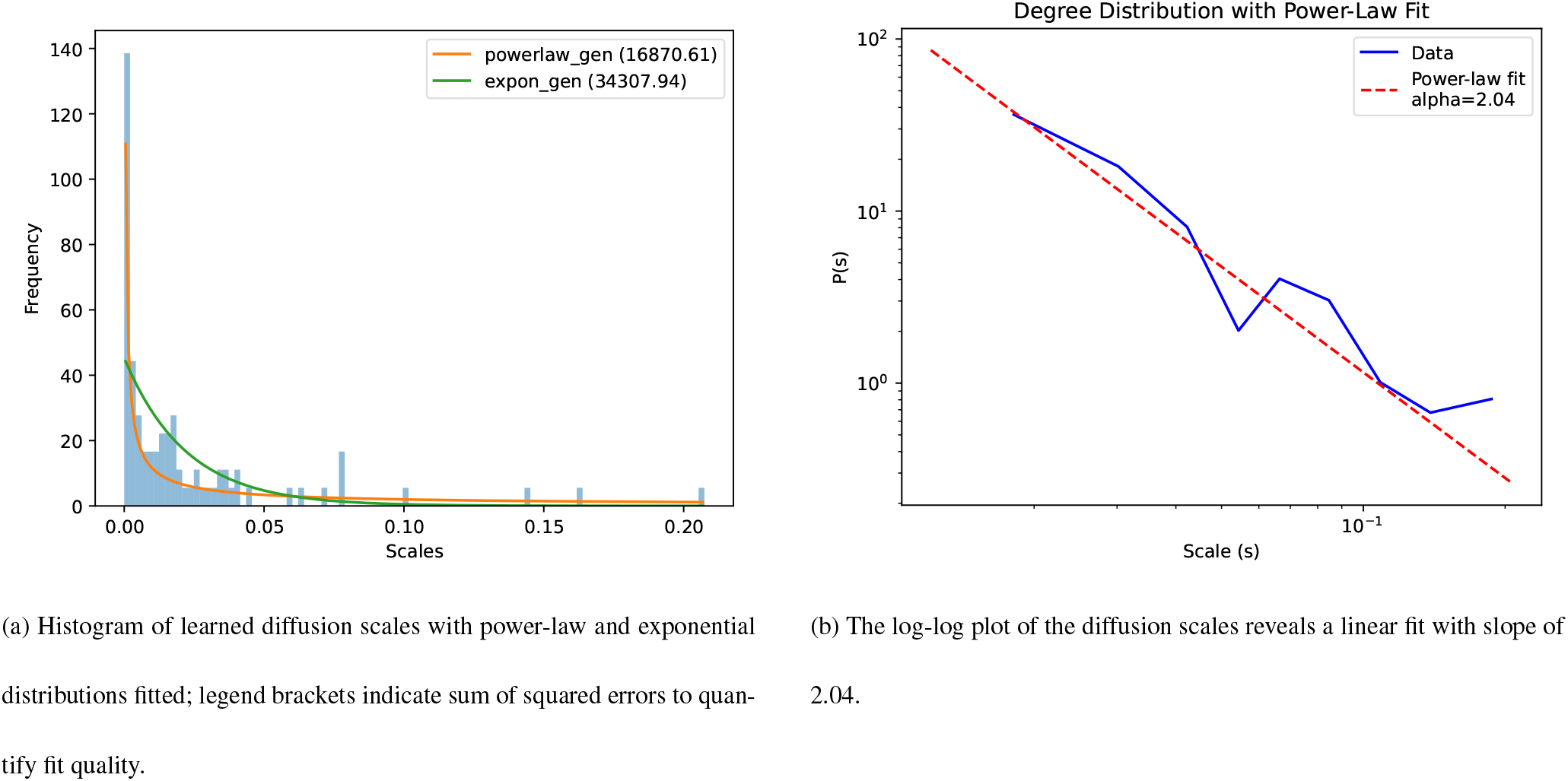
Comparison of distribution of diffusion scales against theoretical distributions.

### Computational Efficiency

For computational efficiency comparisons, we chose A-GHN model as it is the best model that combines graph diffusion, heat kernels, and graph neural networks. We compare our model with A-GHN across various factors such as the number of parameters, model size and the time taken to run on GPU. Both assume that the Laplacian spectrum is pre-computed. A-GHN combined multiple diffusion scales using attention which has an implicit quadratic time complexity. Whereas, in the proposed model we learn the fundamental diffusion scales and the linear layer dramatically reduces the time required for training (see Table 3). We see from the results that while the performance results are superior for the proposed model, it is accomplished with a model that is 4 times smaller in size (number of parameters) and trains at double the speed as compared to A-GHN. More importantly, the proposed model does not compromise on the interpretability aspect. The interpretability afforded by A-GHN arises from the attention weights reflecting the importance of manually preset diffusion scales for predicting FC. On the other hand, in the proposed model, the diffusion scales are not predefined but learned and allow us to interpret region-wise differences in the scales of diffusion.

**Table 3:** Computational requirements of our model compared to the previous state-of-the-art (SoTA) model, A-GHN. Our model shows significant improvements over A-GHN across factors such as model size and computational time.

| Model | Num Parameters | Model Size | GPU Time (s) |
| --- | --- | --- | --- |
| Graph Wavelet | **61,248** | **0.058 MB** | **7.13** |
| A-GHN | 244,648 | 0.233 MB | 12.88 |

## DISCUSSION

In this paper, we used the graph diffusion wavelets to perform multi-scale and multi-resolution analysis of the brain connectome. This method predicts the spatial extent to which the RoIs communicate in terms of the diffusion scales that range from local (fine/small communities) to global (course/large communities) scales. Instead of searching for the correct (optimal) scale by trial and error, the proposed framework allows for learning the diffusion scales using a downstream task of mapping to corresponding static resting-state FC. Our contributions are 2-fold: (1) We learn the SC-FC mapping with state-of-the-art results quantified by Pearson Correlation (0.8326) using an end-to-end linear and explainable graph wavelet diffusion, and (2) The learned diffusion scales follow a power-law distribution conforming to the well-established scale-free property of brain networks. The results suggest that the learning process of diffusion scales enables determination of neighborhood structure appropriate for estimation of FC from SC.

### Learning Node-Specific Diffusion Scales

In the following we explain how the method of graph diffusion wavelet and its application on the human structural connectome for the purpose of deriving the FC leads to tuning of RoI-specific diffusion scales. The graph diffusion wavelet is a probing mechanism which applies an impulse individually at each RoI and observes the fashion in which the impulse travels through the rest of the connectome. In other words, the wavelet captures the response of the connectome to an impulse applied at a specific RoI. Now, each RoI can communicate with other RoIs and the spatial extent of this communication is decided by the magnitude of diffusion scale. If the diffusion scale is smaller, then the RoI forms a local communication network and if the diffusion scale is larger, then the RoI forms a global communication network. Since the dynamics related to function reside on the structure, the diffusion scales in the model are dependent not only on FC but also on SC. Thus, the correct way to obtain the scales is by allowing diffusion at individual nodes at different scales in such a way that it gives rise to FC, unveiling the functional aspects of the individual RoIs. The localization aspect of wavelets helps in identifying the optimal scales at which the individual RoIs operate, indirectly identifying the size of communication networks regulated by the RoI.

### Structural similarity may not correspond to functional similarity

The diffusion scales as a measure of size of communication network is not the only aspect of graph diffusion wavelets. Wavelet brings about localization of the diffusion and each wavelet is characterized by a scale which depends on the FC. We initialize all the scales to 0 and thus the scales will only increase in response to the activity caused by the underlying task or, as in this case, the task of mapping SC to resting-state FC. It can be proved that two structurally identical nodes will have similar wavelet representation at the same scale (Donnat et al., 2018). But due to our downstream task, even if any two RoIs are structurally similar, different functional roles nudge them to have different scale and thus different wavelet representation. In other words, functional roles of the RoIs dictate the spatial extent to which they communicate regardless of having similar neighborhoods. Such interpretations of diffusion scale reveals an interesting mechanism for information transfer in the brain. The diffusion scales are learned by the model only for the objective of prediction of FC, which enables it to encode both the structural aspect of a node and functional pattern observed in the brain.

### Adaptive selection of Diffusion Scales

Another architectural advancement over the previous methods is that the previous methods manually selected the diffusion scales for the single or multiple heat kernel models (Abdelnour et al., 2014; Oota et al., 2024; Surampudi et al., 2018). Since the scale essentially captures the relationship between structure and function, it would be appropriate for the model to learn the scales instead of manual selection. The proposed framework fills this gap of choosing the diffusion scales by learning them using backpropagation of error during training. The diffusion scale update rule can be mathematically represented by a closed form equation, leading to an iterative update scheme. Since these scales are learned in a data-driven manner, they capture the cohort-level trend of diffusion scales.

### Diffusion Scales and Scale-free Property

Third contribution based on our results is that the brain network exhibits a scale-free architecture, characterized by the distribution of diffusion scales across its nodes. Our observations reveal that these diffusion scales adhere to a power-law distribution, indicating that a minority of nodes interact with a large number of nodes, while the majority interact with only a few. Diffusion scales serve as a proxy for degree distribution; if a network’s degree distribution follows a power-law, it may potentially be a scale-free system. The presence of scale-free properties in brain connectomes has been previously documented (Bassett & Bullmore, 2006; Eguíluz, Chialvo, Cecchi, Baliki, & Apkarian, 2005; Sporns & Kötter, 2004). This scale-free organization is crucial for optimal brain function and suggests that the brain operates at a critical point (Beggs, 2022). Several studies have shown that essential neural systems enhance the transmission, storage, and processing of information through this architecture (Marinazzo et al., 2014; Shew, Yang, Yu, Roy, & Plenz, 2011).

### RoI-specific Diffusion Scales and Resting State Networks

Based on learned diffusion scales we identified that frontal pole has the largest community structure. The frontal pole of the brain plays a significant role in various resting-state networks (RSNs), particularly through its involvement in the default mode network (DMN), the frontoparietal control network (FPN), and the social emotion network (SEN) (H. Liu et al., 2013). Other brain regions forming hubs mostly belong to the temporal lobe, which includes Banks of the Superior Temporal Sulcus, Transverse Temporal and Left Parahippocampus. Another region belonging to occipital lobe which has large diffusion scale is pericalcarine region. Although these regions are not necessarily related to default mode network, they are involved in other sensory/cognitive processes such as auditory, visual, social, and memory tasks.

### Scalability and Sensitivity to Noise

Consider the scalability of diffusion wavelets. To analyze this, refer to Table 3 that compares the number of learned parameters with those of A-GHN. Our model learns *N* diffusion kernels and the weights of linear layer are learned in a computationally efficient manner, resulting in a training time of 7.13 seconds for 100 epochs. A potential bottleneck is the 87 matrix multiplications required for heat kernel calculation, but this can be parallelized with minimal GPU requirements. Alternatively, Chebyshev polynomials can approximate the heat kernel to increase speed (Hammond et al., 2011). Compared to A-GHN, which employs multiple GCNs and an attention mechanism, our method is more computationally efficient. Additionally, the proposed model is more efficient in both time and space complexity than MKL that uses LASSO optimization algorithm for combining heat kernels.

Wavelet methods are recognized for their robustness to noise in the signal processing domain, where they can regularize noisy data, such as in point clouds (Hammond et al., 2011). Donnat et al. demonstrated that the difference in wavelets of nodes with equivalent neighborhoods in a noisy graph has an upper bound that depends linearly on the amount of perturbation introduced in the graph (Donnat et al., 2018). In our perturbation studies (see supplementary material), we observe that when the model is trained on perturbed SC, the wavelets resist the added noise, resulting in FC reconstruction with approximately 70% Pearson’s correlation for some test subjects.

## CONCLUSION

Overall, we have developed a method which leverages the graph wavelets in order to obtain explainable and efficient brain structure-to-function mapping through the multi-scale, multi-resolution properties. We compute a wavelet for each node operating at a unique scale and learn the node-appropriate scales during the training process. We overcome the problem of manually choosing the diffusion scales by creating an end-to-end function for efficiently and accurately modelling the relationship. The resulting model has fewer number of trainable parameters and also finds meaningful parameters associated with the brain regions. Our model not only captures the SC-FC mapping but also identifies the community structure supported by individual RoI. In future, this study could be extended to task-based fMRI data as well as for resting state fMRI data related to ageing and neurodegenerative diseases. We expect that the diffusion scales learned in such cases could potentially be used as biomarkers.

## ACKNOWLEDGMENTS

The authors thank Dr. Subba Reddy Oota, INRIA, Bordeaux, France for sharing the code and giving guidance related to dataset preparation which proved to be very helpful for conducting various experiments.

